# Pyrophilic smoke flies (Diptera: Platypezidae: Microsaniinae) from prescribed burnings in boreal Finland: occurrence and behaviour of adult flies and first observations of immature stages

**DOI:** 10.64898/2026.08.31.748078

**Authors:** Tapani Neuvonen, Victoria G. Twort, Gunilla Ståhls

## Abstract

Smoke flies (Diptera: Platypezidae: *Microsania*) are among the most specialized pyrophilic insects, exhibiting a strong and rapid reaction to smoke. Although their attraction to smoke has been known for a century worldwide, their immature stages and breeding habitat remained undiscovered. We hand-netted smoke flies at seven prescribed forest burns in boreal Finland between 2022 and 2024 and took soil samples in 2026. Smoke fly swarming and reproductive behaviour was observed at all burn sites. We obtained smoke flies of five out of the six Finnish species, with *Microsania pallipes* dominating (94%). Following the behavioural observations at burn sites, we later screened soil samples and found larvae identified as *Microsania* mtDNA COI barcodes in the burnt soil. We provide a first description of the larva of *Microsania straeleni* (Microsaniinae). Our discovery of the larval habitat in burnt soil resolved the long-standing knowledge gap on smoke fly breeding sites. The morphological characteristics of the immature stages will help elucidate the phylogenetic placement of the Microsaniinae within the Platypezoidea lineage.

## Introduction

Fire is a fundamental ecological force. The earliest evidence of naturally occurring wildfires dates to over 400 million years ago (Glasspool et al., 2004). For animals the post-fire habitats offer immediate and long-lasting benefits, including increased availability of nutrients and the removal of existing competitors. Over millions of years many organisms have evolved to exploit these conditions and, to a varying extent, have become dependent on them.

Among organisms most directly affected by changes in natural fire regimes are pyrophilic insects, species characterized by a close association with wildfires and post-burn habitats (for a recent review see Bell, 2023). These insects are attracted to fire or smoke and have reproductive association with recently burned environments. Pyrophily has evolved independently multiple times across several insect orders, including Coleoptera, Diptera, Hemiptera, Hymenoptera, and Lepidoptera, with over 200 species classified as pyrophilic globally. Pyrophilic insects are typically among the earliest colonizers of burnt areas, some taxa arriving while the fire is still active, but are relatively rare or completely absent in unburnt habitats (Bell, 2023).

The smoke flies, the genus *Microsania* Zetterstedt (Diptera: Platypezidae: Microsaniinae), are small flies with a body length of 2.5-4 mm. *Microsania* is the single genus in the subfamily Microsaniinae and is a truly cosmopolitan member of Platypezidae distributed across all biogeographical regions except Antarctica, with 22 described species globally (Chandler, 2001). In Finland six species of *Microsania* are recognized (Ståhls, 2014).

Among the platypezids, *Microsania* is distinguished by its fumotropic behaviour, being attracted to smoke, and found nearly exclusively in association with fire and smoke. This behaviour is reflected in the vernacular name, the smoke flies. It was first reported by Severin (1921) in Europe and since documented globally, including from Africa (Collart, 1936a), Australia and New Zealand (Collart, 1934a), northern Europe (Chandler, 2001; Ståhls & Rättel, 2013), North America (Kessel, 1947; Malloch, 1935; Snoddy & Tippins, 1968), and the Pacific island of Fiji (B. Sinclair & Chandler, 2007). All 22 globally described species exhibit this behaviour and are associated with wildfires. Upon arriving at a smoke source, *Microsania* forms aerial swarms inside the smoke plumes. Swarms can consist of hundreds of individuals, closely tracking the smoke plume as it shifts with wind (Bickel, 1996; Klocke et al., 2011; Snoddy & Tippins, 1968). The swarms typically consist mainly of males, which then attracts the females and provides an opportunity for mating (Bickel, 1996; Chandler, 1978; Snoddy & Tippins, 1968). While this behaviour is well documented, the adult habitat of any *Microsania* species outside of fires was never found, except for single observations of adult flies (e.g. Collin, 1934; Ebejer & Andrade, 2010; Melander, 1922; Wikars, 1997). Any potential larval habitat based on the adult behaviour observations remained unidentified. Several larval habitats have been suggested, including fire-associated fungi, forest soil and burnt wood or soil (Collart, 1934, 1936a, 1958; Kessel & Maggioncalda, 1968; Klocke et al., 2011; Tkoč et al., 2017; Wikars, 2009), but attempts to locate or rear the larva have been unsuccessful (Collart, 1936b; Janssens, 1949) and the immature stages have remained completely unknown (Chandler, 2001; B. J. Sinclair & Cumming, 2006).

Another poorly understood aspect of smoke fly biology is their association with phoretic mites. Phoretic mites are frequently observed attached in clusters to the ventral surface of the abdomen in *Microsania*, and already Kessel (1947) noted a group of up to 80 mites on a single fly. Phoretic mites are not documented in other Platypezidae taxa (Chandler, 2001). As phoretic mites typically share the reproductive habitat of their host and rely on the host for dispersal between suitable substrates (Walter & Proctor, 2013), identity and ecology of the mites carried by *Microsania* has been suggested as a route to uncovering the unknown larval habitat of the genus (e.g. Chandler, 2001; Collart, 1933; Edwards, 1934; Janssens, 1949; Kessel, 1947). However, the ecological significance of this association and the timing of mite attachment and whether it was in above-or below ground terrestrial environment remained unexplored.

Although smoke flies can occur in very large numbers in recently burned areas, they disappear from burned sites within days (Chandler, 2001). The lack of research on smoke flies can be partly explained by the practical challenges of studying pyrophilic insects: forest fires are typically locally unpredictable events, and observations within days of a wildfire require coordination with fire management authorities. Most literature on the genus relies on observations made at small scale fires, such as bonfires. While this method can attract smoke flies for opportunistic collecting, no studies have attempted to systematically map the species assemblages of the flies at large scale fires.

This study examines smoke flies found at prescribed burns within one day of the onset of the fire. Specifically, we document the occurrence and behaviour of *Microsania* at seven sites of prescribed burns sampled between 2022 and 2024 and in 2026 specifically to investigate possible the larval habitat. Our objectives are to: (i) describe and compare the *Microsania* species assemblages present at prescribed burns in Finland; (ii) document behaviours under active fire conditions, including swarming, mating, and oviposition; (iii) find and identify smoke fly larval habitat and immature stages and (iv) characterize the phoretic mites associated with Finnish smoke fly populations.

## Material and methods

### Study sites

Field observations and specimen collecting was conducted at seven prescribed burning sites in southern and northern Finland between August 2022 and June 2024 (Table 1). All burns were conducted by Metsähallitus (Finnish Forest Service) as part of conservation restoration or silvicultural management on state-owned lands. Prescribed burning is a forest management practice used for forestry or agricultural and biodiversity promoting purposes. In Finland, it is used routinely to combat the declining amounts of natural fires (Lindberg et al., 2020) that has led to a lack of post-burn habitats contributing to the decline of more than 30 species of mainly Coleoptera (Hyvärinen et al., 2019). Approximately 400 ha of forested area is burned annually in prescribed burnings nationwide (Koivula & Vanha-Majamaa, 2020; Lindberg et al., 2020; Vanha-Majamaa et al., 2007). Visited sites were selected based on the availability of burns within the burn schedule of Metsähallitus.

**Table 1.** Summary of prescribed burning sampling sites. RB=restoration burn, SC=silvicultural nature management burn after clearcut.

| Site | Burn type | Nature type | Coordinates<br>(Lat °N, Lon °E) | Burn date | Area<br>(ha) |
| --- | --- | --- | --- | --- | --- |
| Evo | RB | herb-rich heath forest | 61.22026,<br>25.20074 | 15 August 2022 | 7 |
| Loppi | RB | sub-xeric heath forest | 60.67546,<br>24.29781 | 22 May 2023 | 5 |
| Nuuksio 23 | RB | barren heath forest | 60.33931,<br>24.56118 | 13 June 2023 | 6 |
| Nuuksio 24 | RB | herb-rich heath forest | 60.33830,<br>24.50691 | 22 May 2024 | 6 |
| Posio | SC | mesic heath forest | 66.03028,<br>27.64670 | 27 May 2024 | 20 |
| Kemijärvi | SC | mesic heath forest | 66.838500,<br>27.257980 | 30 May 2024 | 30 |
| Sodankylä | SC | sub-xeric heath forest | 67.63652,<br>27.26668 | 6 June 2024 | 20 |

Sites spanned approximately 800 km in the north–south direction at 90–230 m above sea level within the boreal climatic zone (Figure 1). Burned areas ranged from 5 to 30 ha in size. The areas encompassed a range of boreal forest types, from herb-rich to barren heath forests.

**Figure 1.**
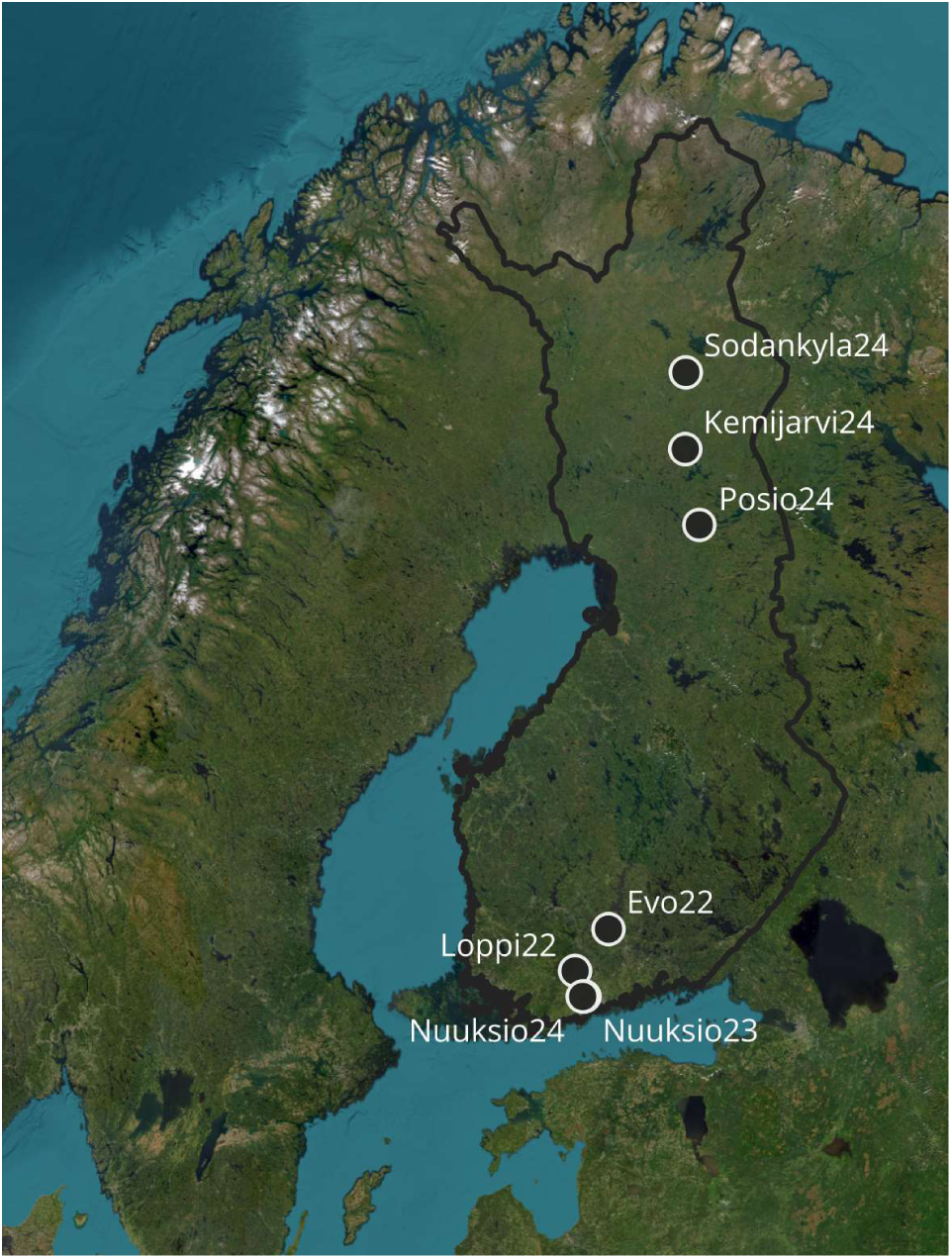
Map of localities of prescribed burns.

Vegetation on the sites was of typical boreal forests, with Scots pine (*Pinus sylvestris*) as the dominant tree species mixed with varying amounts of Norway spruce (*Picea abies*) and birches (*Betula* spp.). Site characteristics, including main forest type, coordinates, burn date, and burned area size are summarized in Table 1.

### Specimen sampling

Burned areas were attended from fire ignition, and sampling commenced once the fire front had passed and smoke-producing areas were accessible, in all cases within two hours of ignition. Specimens were collected by sweep-netting with an entomological net (40 cm diameter) through smoke plumes at approximately 20–180 cm above ground level (Figure 2 A). Sampling covered both burn edges and interior sections of the burned area. The sweep-netting was done by slow walking in the burned areas and focusing on places where the flies could be found swarming at concentrated sources of smoke, such as smouldering tree stumps and fallen trees. When swarming was encountered, behavioural observations were noted before sampling continued. While sweeping, the net was frequently checked for flies, and all caught specimens were immediately transferred into vials filled with 75% ethanol.

**Figure 2.**
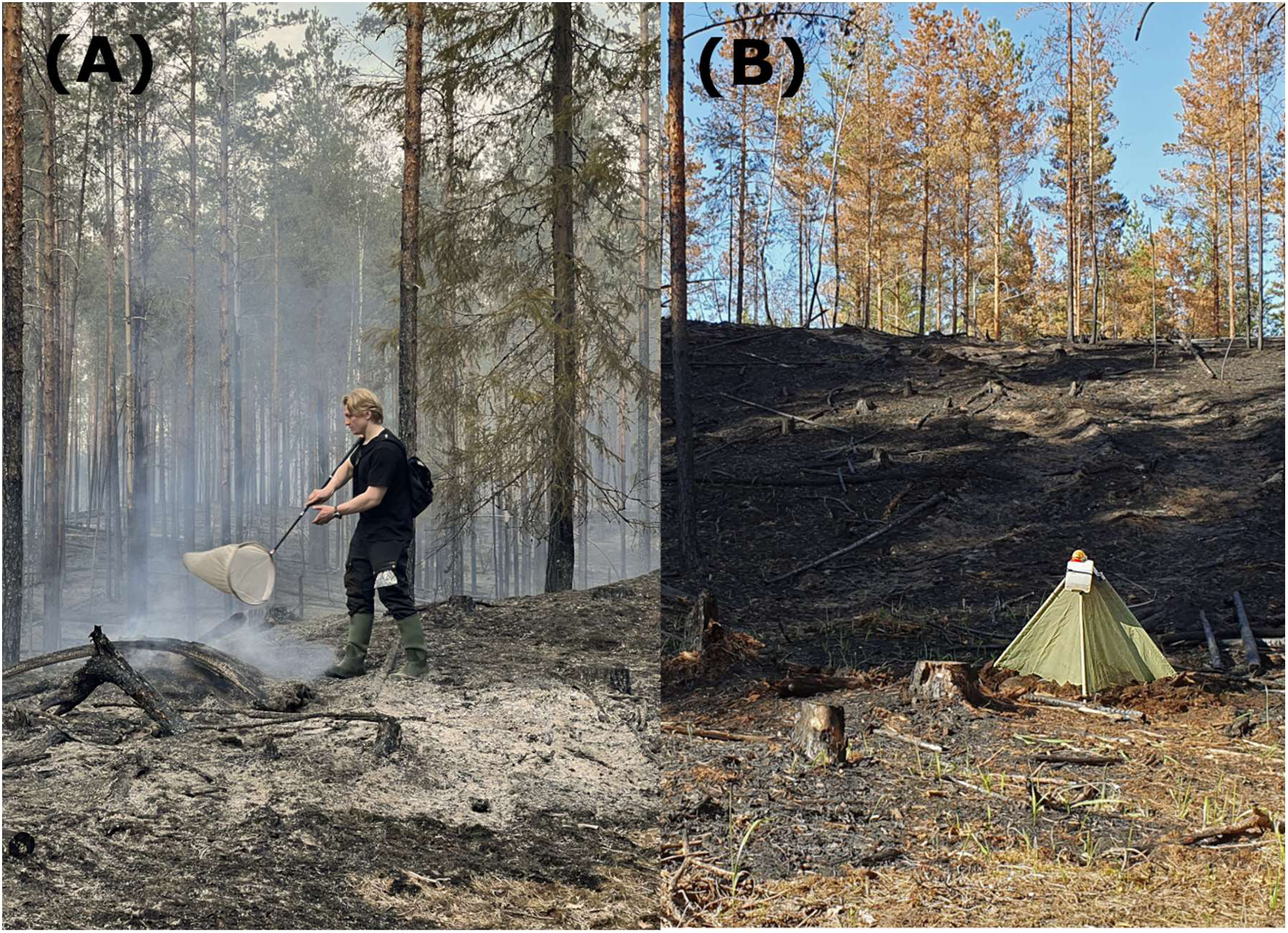
A. Collecting. B. Emergence trap.

#### Sample identification

Collected smoke fly specimens were morphologically identified to species level using the identification keys of Chandler (2001) under a Leica S9E stereomicroscope at the Finnish Museum of Natural History (FMNH) (Helsinki, Finland). The presence of phoretic mites was recorded for each identified specimen. Selected mites were removed and sent for morphological identification to mite expert Dr. Veikko Huhta, University of Jyväskylä, Finland. Mites were also removed and placed individually in tubes in ethanol for later DNA barcoding.

### Emergence trapping and Soil Samples

At the Loppi burn (22 May 2023), female *Microsania* were observed walking on the burned soil where the fire had passed approximately 4 hours earlier and entering small ground crevices near swarm locations. The exact location was recorded as the behaviour was expected to be related to oviposition. The site was revisited 20 days after the prescribed burning (11 June 2023), and an emergence trap was set up in the same location. The emergence trap consisted of a pyramid-shaped steel frame (60 x 60 x 60 cm) covered with thin nylon mesh (Figure 2 B). A flexible tube connected to a plastic container was placed on the top of the trap, and the container was filled with 98% ethanol. The edges of the nylon mesh were sealed to the ground with soil to ensure that only insects emerging from beneath the trap could enter the container. The emergence trap was retrieved after 26 days (7 July 2023), covering days 20-46 after the burn. After collecting the trap, all specimens were identified to species at FMNH, and their phoretic mite presence recorded, as described above.

At a selected prescribed burn in southern Finland (Vihti, 3 June 2026, sub-xeric heath forest, Scots pine dominated, 0,22 ha), female *Microsania* were again observed walking on the burned soil near swarm locations. To investigate whether larvae develop in burned substrates, five samples of soil and litter from the top 5-8 cm layer of the forest floor were collected with a trowel and placed in separate 10 L plastic ziplock bags. The samples comprised of typical humus layer characteristic of sub-xeric heath forest. The fire had passed the area approximately 5 hours before the samples were taken. The temperature of the soil surface ranged between 30-46 °C with hot spots reaching 200 °C (measured with a Extech IR400 infrared thermometer). The bags were stored at room temperature away from direct sunlight and opened daily for airing and inspection of any emerged adults.

### Larval examination and description

The soil samples were screened under a stereomicroscope under low magnification, and any observed immature stages were retrieved and placed individually in 96% ethanol. Larva assigned for DNA barcoding were placed in a conventional freezer in ethanol. A third instar larva (L3) was placed in glycerol and studied and photographed using a Keyence VHX-7000 automated digital high-resolution microscope at FMNH. The terminology used for larval description follows Chandler (2001).

### DNA barcoding

The Phire™ Tissue Direct PCR master Mix #F-170S (Thermo Scientific Baltics UAB, Vilnius, Lithuania) was used for generating DNA barcodes as it is designed to perform PCR directly from tissue samples with no prior DNA purification. DNA was extracted from an entire larva, following the Dilution & Storage protocol with the following modifications: 1) incubated at room temperature for about 20 min and 2) 2 µl supernatant was used in the PCR reaction. The mtDNA COI barcode was PCR amplified using universal primers LCO1490 and HCO2198 (Folmer et al., 1994). Amplified PCR products were electrophoresed on 1.5% agarose gels and treated with Exo-SapIT (USB Affymetrix, Ohio, USA) prior to sequencing. The PCR primers were used for sequencing, which was outsourced to the Sequencing Service Laboratory of the Institute for Molecular Medicine Finland (www.fimm.fi). The sequences were edited for base-calling errors and assembled using Sequencer™ (version 5.0) (Gene Codes Corporation, Ann Arbor, MI, USA) and obtained barcode sequences were submitted to GenBank (accession numbers xxxxxx). The resulting barcodes were combined with adult representatives from all Finnish species and *Megaselia abdita* (Phoridae) as an outgroup. The resulting dataset was analysed using the Neighbor-Joining method (Saitou & Nei, 1987) under the Kimura 2-parameter substitution model (Kimura, 1980), with support being assessed with 500 bootstraps using the software MEGA11 (Tamura et al., 2021). All positions with less than 95% site coverage were eliminated, i.e., fewer than 5% alignment gaps, missing data, and ambiguous bases were allowed at any position (partial deletion option).

## Results

### Species composition and sex ratios

A total of 1,637 specimens of *Microsania* were collected across the seven prescribed burning sites (Table 2). Five of the six species occurring in Finland (Ståhls, 2014) were recorded: *Microsania pallipes* (Meigen, 1830), *M. pectipennis* (Meigen, 1830), *Microsania capnophila* Shatalkin, 1985, *M. vrydaghi* Collart, 1954 and *Microsania straeleni* Collart, 1954. *Microsania pallipes* was the dominant species at all sampled sites, accounting for 93.5% of all specimens (Table 2). *Microsania pectipennis* was the second most frequent species and was present at six sites. *M. capnophila* was found at five sites in low numbers, and *M. vrydaghi* was collected at four sites at similarly low numbers. *M. straeleni* was found at two sites, with only one specimen collected at each site.

**Table 2.**
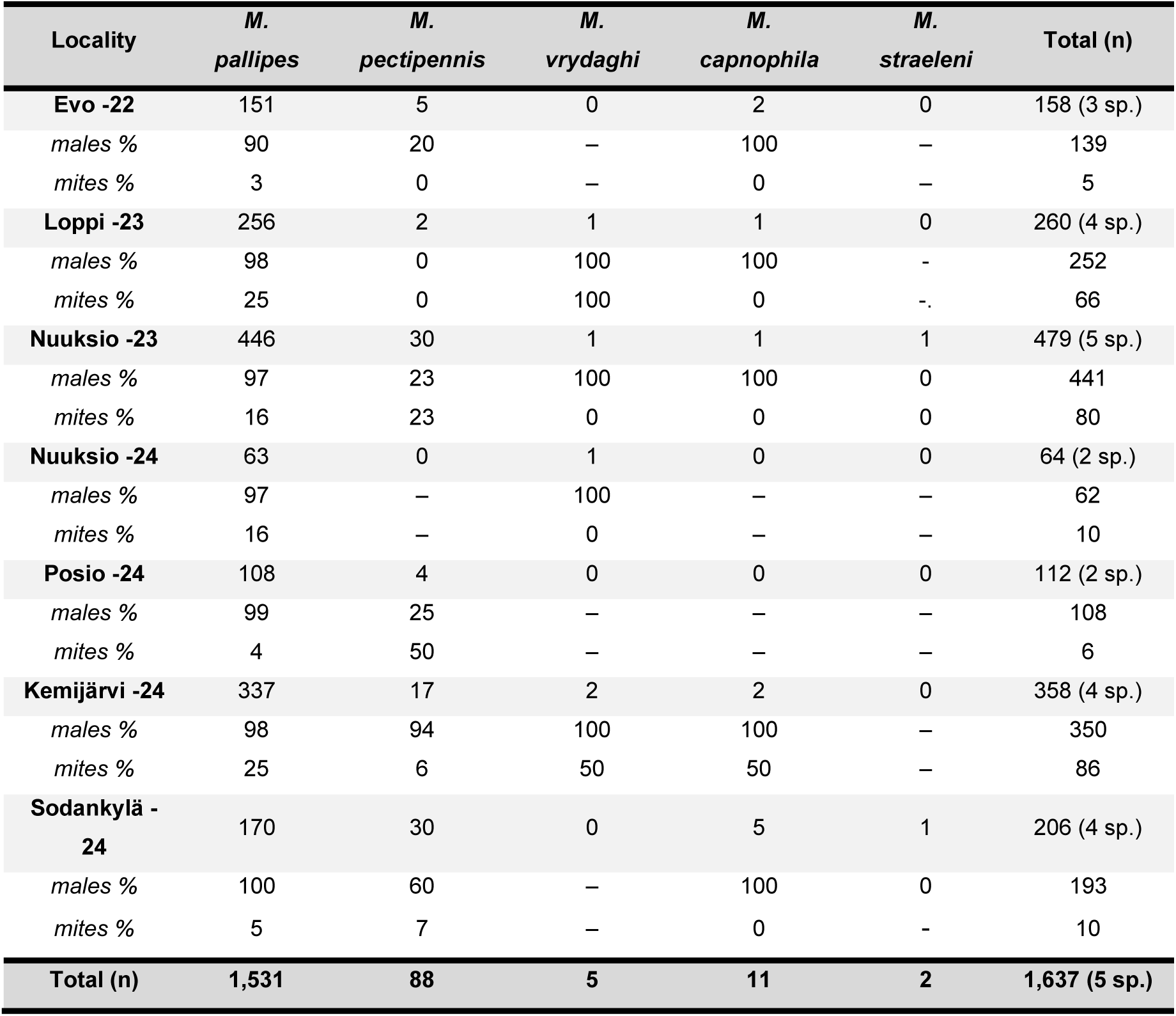
Microsania species collected at each burn site with specimen numbers. The last column indicates the number of specimens for each row.

| Locality | *M.*<br>*pallipes* | *M.*<br>*pectipennis* | *M.*<br>*vrydaghi* | *M.*<br>*capnophila* | *M.*<br>*straeleni* | Total (n) |
| --- | --- | --- | --- | --- | --- | --- |
| **Evo -22** | 151 | 5 | 0 | 2 | 0 | 158 (3 sp.) |
| males % | 90 | 20 | – | 100 | – | 139 |
| mites % | 3 | 0 | – | 0 | – | 5 |
| **Loppi -23** | 256 | 2 | 1 | 1 | 0 | 260 (4 sp.) |
| males % | 98 | 0 | 100 | 100 | - | 252 |
| mites % | 25 | 0 | 100 | 0 | -. | 66 |
| **Nuukio -23** | 446 | 30 | 1 | 1 | 1 | 479 (5 sp.) |
| males % | 97 | 23 | 100 | 100 | 0 | 441 |
| mites % | 16 | 23 | 0 | 0 | 0 | 80 |
| **Nuukio -24** | 63 | 0 | 1 | 0 | 0 | 64 (2 sp.) |
| males % | 97 | – | 100 | – | – | 62 |
| mites % | 16 | – | 0 | – | – | 10 |
| **Posio -24** | 108 | 4 | 0 | 0 | 0 | 112 (2 sp.) |
| males % | 99 | 25 | – | – | – | 108 |
| mites % | 4 | 50 | – | – | – | 6 |
| **Kemijärvi -24** | 337 | 17 | 2 | 2 | 0 | 358 (4 sp.) |
| males % | 98 | 94 | 100 | 100 | – | 350 |
| mites % | 25 | 6 | 50 | 50 | – | 86 |
| **Sodankylä -**<br>**24** | 170 | 30 | 0 | 5 | 1 | 206 (4 sp.) |
| males % | 100 | 60 | – | 100 | 0 | 193 |
| mites % | 5 | 7 | – | 0 | - | 10 |
| **Total (n)** | **1,531** | **88** | **5** | **11** | **2** | **1,637 (5 sp.)** |

No clear trends were found between the characteristics of the sampling sites and the collected specimens. The number of species and the total number of specimens caught varied considerably among sites, with no clear relationship to latitude, time of the year, burn size, or burn type. Between two and five species were recorded at each site, with more species generally found when the number of specimens collected was larger. The overall sex ratio was strongly male biased at all sites, and males accounted for 94% of all collected smoke flies (Table 3).

**Table 3.** Abundance, sex ratio and phoretic mite prevalence for each *Microsania* species (final numbers).

| Species | % of catch | N | % male | Mite prev. % |
| --- | --- | --- | --- | --- |
| *M. pallipes* | 93.5 | 1,531 | 97.1 % | 16.2 |
| *M. pectipennis* | 5.4 | 88 | 48.9 % | 13.6 |
| *M. capnophila* | 0.7 | 11 | 100 % | 9.1 |
| *M. vrydaghi* | 0.3 | 5 | 100 % | 40.0 |
| *M. straeleni* | 0.1 | 2 | 0 % | 0 |
| Total | 100 | 1,637 | 94.4 | 16.1 |

### Phoretic mites

Overall, 16.2% of all *Microsania* specimens collected by hand netting carried phoretic mites at the time of examination (Table 3). This does not account for mites that could have detached during handling of fly specimens, as mites were frequently seen loose in the vials. Among net-caught flies, females carried mites about half as often as males (23.0% vs. 15.7%). The numbers of phoretic mites attached to individuals varied greatly. Some specimens had one or two mites, while others carried a significant load attached to the ventral side of their abdomen. In an extreme occasion, a single *M. pallipes* male collected from Evo was carrying 87 mites attached to the abdomen.

Mites were most frequently attached to the ventral surface of the basal three abdominal segments, arranged in rows or aggregated in a dense cluster. Singular mites were also found attached to the antennae, coxae and male genitalia. Individual flies occasionally carried mites belonging to more than one taxon.

Based on morphological identification, adult mites were identified as *Pediculaster mesembrinae* (Prostigmata: Pygmephoridae) and nymphal mites as deutonymphs of *Dendrolaelaps* sp. (Mesostigmata: Digamasellidae). Further morphological identification of the deutonymphs to species level was not possible due to lack of the key morphological characteristics for identification are expressed only in the adults (V. Huhta, pers. comm.).

### Observations of smoke fly behaviour

Swarming in smoke was observed at all sites within the first three hours after fire ignition. The swarms consisted mainly of males (Table 2), with swarm size varying from tens of individuals to >100. The swarms were observed where visible smoke was present, including at smouldering tree stubs or fallen trunks, or at other localized smoke sources. Within the smoke plumes, *Microsania* formed roughly elliptical swarms approximately 50-180 cm above ground and as estimated typically within 3 meters from the smoke source. The swarms tracked the smoke plume closely as it moved according to the wind direction. When part of the swarm was collected by sweep-netting, additional flies quickly moved in to replace those removed. However, if a swarm was collected in its entirety, a new swarm did not form immediately in the same location. Up to three species of *Microsania* were collected from a single swarm, predominantly consisting of the most abundant species found in this study, *M. pallipes* (Table 2).

Copulation was directly observed at several times at two sites (Evo, Sodankylä) near swarming events. In one instance, copulating individuals were observed with both sexes bearing a substantial load of phoretic mites (Figure 3).

**Figure 3.**
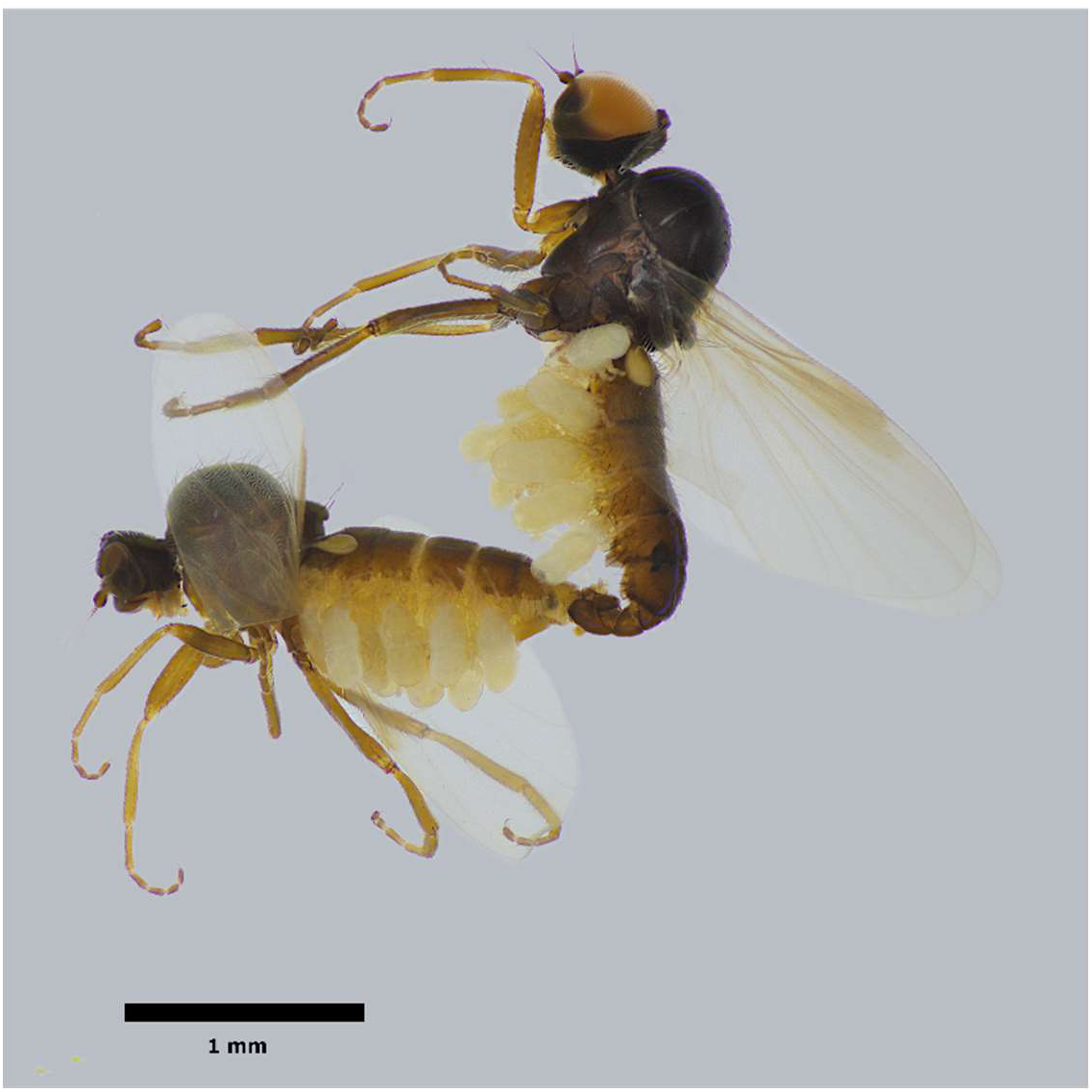
Mating pair of *Microsania pallipes*. with mites attached.

At several localities, females were observed walking rapidly on the burned soil near swarm locations and seen entering small crevices in the burned soil. Males were observed on the soil surface but did not exhibit the same rapid movement or enter the soil.

In burned areas at the first day after the fire, we observed that *Microsania* was the most abundant insect flying in the sampled areas. Only a small number of other insects were caught as bycatches during the hand-netting.

### Emergence trapping and Soil samples

The emergence trap was collected after 26 days of collecting, being deployed from 20 to 46 days post-fire. The trap contained a total of 22 *M. pallipes*, consisting of 4 males and 18 females. All specimens were lightly pigmented with translucent cuticle, indicating that they had entered the trap shortly after emerging.

Following daily monitoring of the soil samples, larvae were seen on 14 June 2026, 11 days after the soil was collected. First adult smoke flies emerged flies emerged from the samples 14 days after collecting, with several more emerging throughout the observation period. The observation was stopped on 4 July when the last adult emerged on a closed petri dish. Larvae were collected for morphological description 20 days after the prescribed burn. In total 10 larvae of different instars were stored in 75% ethanol and examined in glycerol.

### DNA barcoding: larval species identification

A barcode datamatrix was constructed from mtDNA COI barcodes of two larvae and multiple barcodes of representatives of the adult stage of all the six Finnish *Microsania* species. The complete dataset consisted of 630 bp, and 55 ingroup specimens. In the resulting barcoding tree (Figure 4) the barcodes of the two larvae showed highest similarity to and were referred to the same clade as included barcodes of adult flies of *Microsania straeleni* Collart.

**Figure 4.**
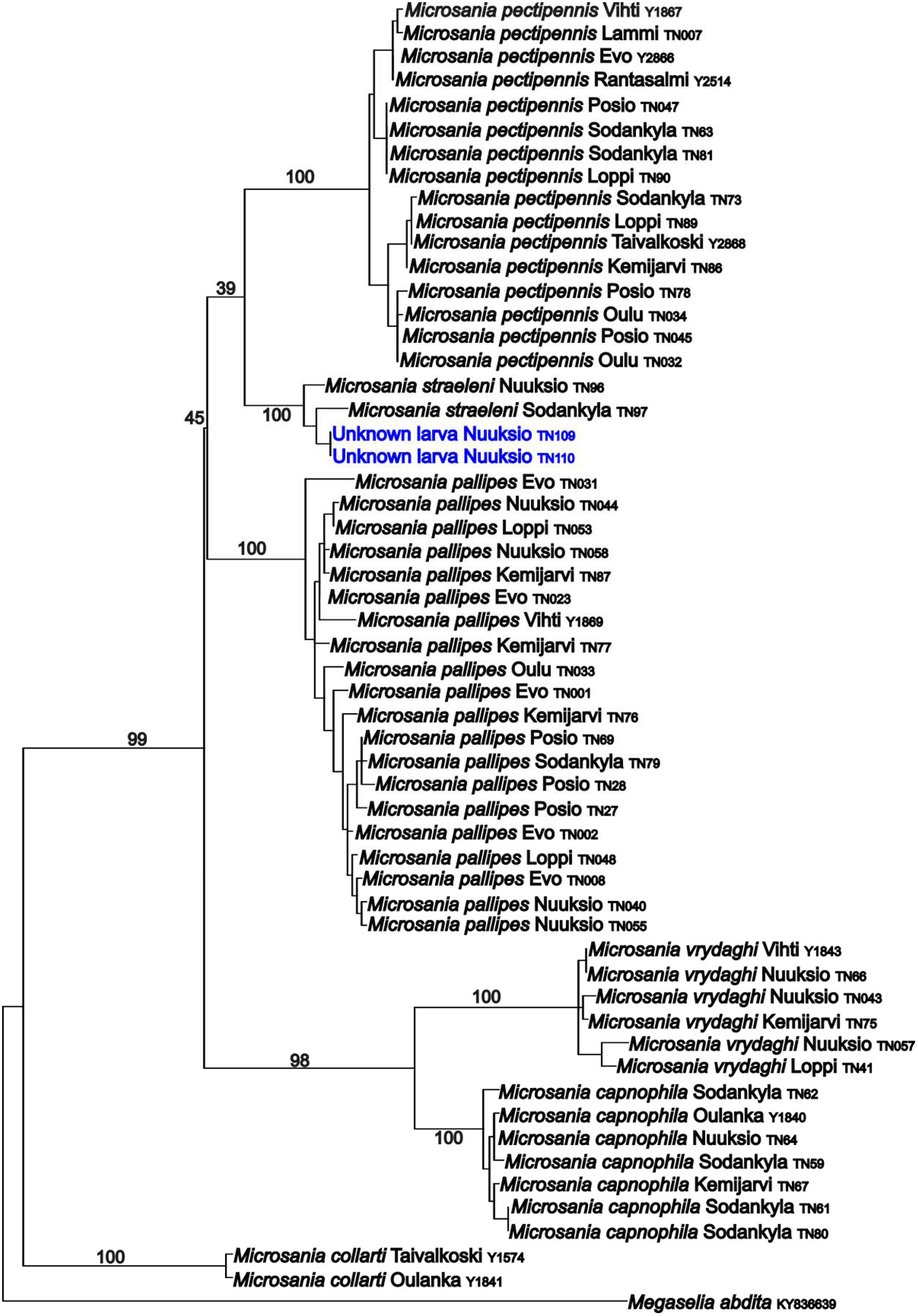
Neighbour-joining barcoding tree for identification of larvae. Species names are followed by collection locality and sample code. Larval samples are in blue. Bootstrap support values are provided only for the backbone topology and major branching events; intraspecific node support values are omitted for clarity.

### Preliminary description of larva of Microsania straeleni

#### Overall appearance

A subcylindrical larva, somewhat dorsoventrally depressed, tapering anteriorly (Figure 5). Marginal and dorsal projections vary in length. Colour ranges from pale yellow to light brown. Body length from anterior margin of prothorax to posterior margin of anal segment approximately 2 mm. All body processes conical with short hair-like structures (setulae) set in a ring-like fashion. Dorsal processes from mesothoracic to last abdominal segment approximately of equal length, while subdorsal processes are shorter. Spinules are present predominantly dorsally on the mesothoracic to the last abdominal segment. The ventral surface of the body is developed into a series of fleshy creeping welts. Segments latero-ventrally with a short process. Thoracic and abdominal segments generally with a total of 10 processes on each segment.

**Figure 5.**
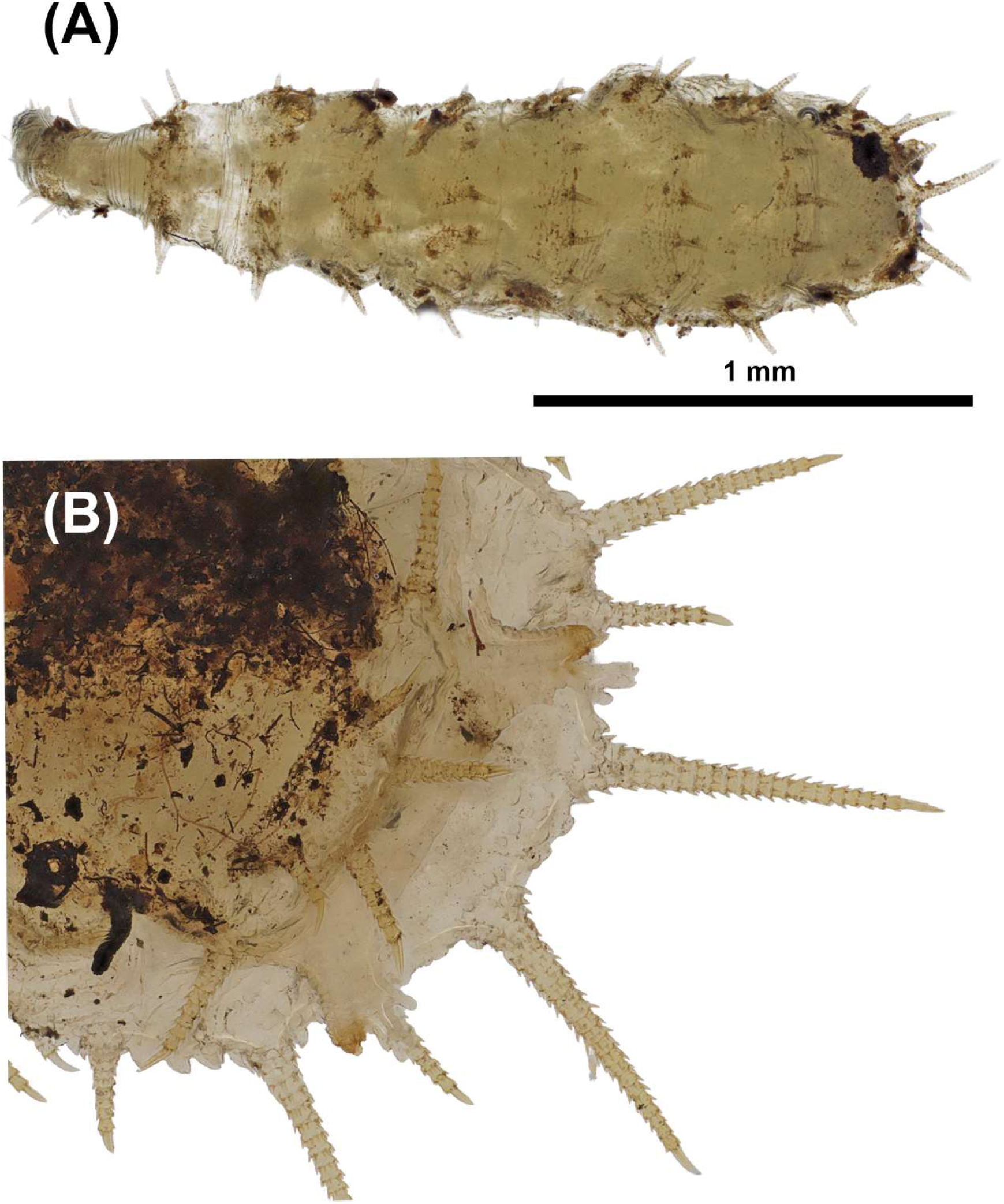
A: Dorsal view of entire larva, B. Ventral view of anal segment.

#### Head

Head and prothorax membranous, small and parallel sided, head narrower than prothorax. Head and prothorax ventrally directed. Antenna long and tapered.

#### Thorax

Mesothorax and metathorax with 6 processes on anterior margin. Mesothorax is narrower than meta-and abdominal thoracic segments.

#### Abdomen

Processes on abdominal segments 1-5 are situated approximately medially on the segment. Processes on the last abdominal segment and anal segment are situated approximately on the posterior margin. Abdominal segments with 2 lateral processes on each side and 6 dorsal processes across the segment, a total of 10 processes on each segment. Anal segment with 8 processes dorsally and 4+2 processes situated ventrally, and with a pair of elaborate posterior spiracles (best visible from ventral side) on intervening area.

## Discussion

### Species composition

Five of the six Finnish *Microsania* species were collected across the seven prescribed burnings. *M. pallipes* was by far the most abundant, accounting for 93.5% of all specimens. This contrasts with reports from other parts of Europe, where *M. pectipennis* has frequently been found as the most common and abundant species at smoke sources (Chandler, 1974; Collart, 1933, 1960; Edwards, 1934). Whether this difference reflects actual biogeographical variation in species abundance or merely differences in sampling methodology and habitat type is unclear. Sampling in the present study was made at large-scale prescribed burning in boreal coniferous forests, while much of the European literature is based on bonfire collecting at temperate deciduous or mixed forests.

The species composition recorded in this study is likely to be biased due to the sampling method. Collecting by sweep netting through smoke plumes may unintentionally focus on certain species and sexes. Superficially the swarms in smoke compose of up to three species of *Microsania*, but each species probably aggregates at specific places in the smoke plume. Each species could possibly occupy the plume at different heights and distances and hand sweeping fails to capture this variation. Species could also differ in their preference of swarming based on distances from the burned forests edge or in different microhabitats within the burn.

While several species appear to be present at the freshly burned areas, these variations could explain the low number of specimens collected for certain species. Similarly, temporal differences in the response to fire could explain the variance in abundance: if species differ in their phenology or at the speed at which they respond to fires, a single day of sampling may overlook species that arrive later. No clear differences in species composition were observed between early-season (May) and late-season (August) burns.

The strong male bias of the collected *Microsania* specimens is congruent with literature observations. Similar sex ratios have been reported from Europe (Chandler, 1974; Collart, 1933, 1933; Edwards, 1934; Severin, 1921), Australia (Bickel, 1996; Collart, 1938) and North America (Snoddy & Tippins, 1968). Edwards (1934) noted that female smoke flies tend to fly very close to the ground. If this is the case, the observed male-female bias could be due to bias resulting from the collecting method. The contrasting sex ratio from the emergence trap (18 females, 4 males) supports this hypothesis. The net-sweeping height may introduce a strong bias against females if they fly near or on the ground surface.

Despite the abundance of smoke flies at prescribed burnings, sometimes with hundreds of individuals collected within hours of ignition, *Microsania* remains remarkably rare in the absence of fire. Very few specimens have been collected outside of fire or smoke globally and smoke flies are not evaluated in the Finnish Red List due to insufficient data (Hyvärinen et al., 2019). Some specimens of *Microsania* have been found in malaise traps at burned areas up to three years after a forest fire (S. Karjalainen, pers. comm., Wikars, 2009), but it is unclear whether the represent actual local populations or occasional dispersing individuals.

### Reproductive behaviour at burn sites

Females were observed walking rapidly on burned soil (T. Neuvonen pers. obs.) and entering crevices in the burned organic layer, consistent with descriptions by Edwards (1934) and Collart (1958). Although oviposition was not directly observed, the discovery of *Microsania straeleni* larvae and puparia from soil samples confirms that females oviposit in the burned substrate. In Australia, *Microsania australis* Collart was observed entering a burnt cavity in a *Eucalyptus* tree trunk (Klocke et al., 2011), which may represent a similar behaviour.

### Mite phoresy

On average, 16% of *Microsania* collected by hand netting carried phoretic mites. The two identified mite taxa, *Pediculaster mesembrinae* and deutonymphs of *Dendrolaelaps* sp., occupy a wide range of habitats. Species of the genus *Dendrolaelaps* inhabit various habitats, including decaying wood, forest soils, ant hills, and bark of fallen logs (Huhta, 2016), and are commonly found phoretic on bark beetles (Pernek et al., 2012) and flies (Mumcuoglu & Braverman, 2010). *Pediculaster mesembrinae* is phoretic on a variety of fly hosts, especially those inhabiting dung and organic waste (Kontschán & Hornok, 2019), and feeds on several types of fungi, particularly moulds (Hernández-Abarca et al., 2005; Szafranek & Lewandowski, 2017).

The relatively high number of phoretic mite carrying *Microsania* from emergence traps (82%) compared to net-caught specimens (16%) provides insight into the timing of mite attachment. It indicates that the mites are present in the soil substrate and attach to the flies at or shortly after emerging. The observation of mating pairs with substantial mite loads suggests that the mites do not prevent successful reproduction, consistent with a non-parasitic phoretic relationship (Figure 3).

## Conclusions

The results presented in this study fills a longstanding knowledge gap regarding the oviposition sites and larval substrate of the smoke flies. In particular, the lack of knowledge of the immature stages of *Microsania* has been identified in multiple worldwide studies since the beginning of the 19^th^ century (e.g. Chandler, 2001; Severin, 1921) and has hampered our understanding of their evolutionary placement relative to Platypezidae (e.g. Tkoč et al., 2017). Here we present for the first time evidence that in the boreal forest environment burned soil is a key factor in smoke fly reproduction. The discovery of *Microsania straeleni* larvae in burn site forest soil constitutes the first confirmed record of the larval habitat of the subfamily Microsaniinae, and the first descriptions of the immature stages of any *Microsania* species.

The preliminary investigation of external morphological characteristics of the *Microsania* larva agree with features such as general shape and the structure and placement of the dorsal and marginal processes on body segments described for other Platypezidae taxa, particularly with those of taxa of Callomyiinae (Chandler, 2001; Ståhls & Twort submitted), but its unique combination of features is not shared with other platypezid taxa.

## Author Contributions

TN: Conceptualization, Methodology, Investigation, Visualization (Image preparation), Writing – original draft, Writing – review & editing. GS: Conceptualization, Methodology, Supervision, Writing – review & editing. VT: Supervision, Visualization (figure preparation), writing – reviewing & editing.

## Acknowledgments

We thank Metsähallitus for providing access to prescribed forest burns, and Veikko Huhta for mite species identification. TN acknowledges financial support for this research from Societas pro Fauna et Flora Fennica, the Environmental Research Foundation of Lammi Biological Station, the Vuokko Foundation for Nature Conservation, the Entomological Society of Finland, Symbioosi ry, and the Kainuu Regional Fund of the Finnish Cultural Foundation.

## Disclosure

The author(s) report no conflicts of interest in this work.

## Notes

### Competing Interest Statement

The authors have declared no competing interest.

